# Isolating neural signatures of conscious speech perception with a no-report sine-wave speech paradigm

**DOI:** 10.1101/2023.11.26.568128

**Authors:** Yunkai Zhu, Charlotte Li, Camille Hendry, James Glass, Enriqueta Canseco-Gonzalez, Michael A. Pitts, Andrew R. Dykstra

**Author notes:** Correspondence: Andrew R. Dykstra or Michael A. Pitts. These authors contributed equally to this work (Enriqueta Canseco-Gonzalez, Michael A. Pitts, Andrew R. Dykstra).

## Abstract

Identifying neural correlates of conscious perception is a fundamental endeavor of cognitive neuroscience. Most studies so far have focused on visual awareness along with trial-by-trial reports of task relevant stimuli, which can confound neural measures of perceptual awareness with post-perceptual processing. Here, we used a three-phase sine-wave speech paradigm that dissociated between conscious speech perception and task relevance while recording EEG in humans of both sexes. Compared to tokens perceived as noise, physically identical sine-wave speech tokens that were perceived as speech elicited a left-lateralized, near-vertex negativity, which we interpret as a phonological version of a perceptual awareness negativity. This response appeared between 200 and 300 ms after token onset and was not present for frequency-flipped control tokens that were never perceived as speech. In contrast, the P3b elicited by task-irrelevant tokens did not significantly differ when the tokens were perceived as speech versus noise, and was only enhanced for tokens that were both perceived as speech *and* relevant to the task. Our results extend the findings from previous studies on visual awareness and speech perception, and suggest that correlates of conscious perception, across types of conscious content, are most likely to be found in mid-latency negative-going brain responses in content-specific sensory areas.

**Significance Statement:** How patterns of brain activity give rise to conscious perception is a fundamental question of cognitive neuroscience. Here, we asked whether markers of conscious speech perception can be separated from task-related confounds. We combined sine-wave speech - a degraded speech signal that is heard as noise by naive individuals but can readily be heard as speech after minimal training - with a no-report paradigm that independently manipulated perception (speech versus non-speech) and task (relevant versus irrelevant). Using this paradigm, we were able to identify a marker of speech perception in mid-latency responses over left frontotemporal EEG channels that was independent of task. Our results demonstrate that the “perceptual awareness negativity” is present for a new type of perceptual content (speech).

## Introduction

How conscious perception emerges from brain activity is a fundamental question in neuroscience (Snyder et al., 2015; Dykstra et al., 2017). Using the so-called contrastive method - contrasting conditions that are equivalent in every respect except for their conscious perceptual contents (Baars, 1988, 2002; Crick and Koch, 1990, 1998, 2003; Dehaene et al., 2003, 2006) - several electroencephalography (EEG) studies have identified (i) an early (∼100-300 ms post-stimulus onset), negative-going response in sensory cortices and a later (> 300 ms), positive-going response near Pz associated with perception. The early response has been termed the perceptual awareness negativity (PAN) (Dembski et al., 2021), and the latter the P3b or “late positivity” (Polich, 2007; Halgren, 2008; Dehaene and Changeux, 2011). Modality-specific versions of the PAN have been recorded in at least three sensory modalities: visual (VAN) (Koivisto and Revonsuo, 2010; Pitts et al., 2012; Shafto and Pitts, 2015; Eklund and Wiens, 2018), auditory (AAN) (Hillyard et al., 1971; Gutschalk et al., 2008; Dykstra and Gutschalk, 2015; Eklund and Wiens, 2019) and somatosensory (SAN) (Jones et al., 2007; Auksztulewicz et al., 2012; Schröder et al., 2019).

One potential confound present in many such studies is the requirement that participants report their percept on a trial-by-trial basis, which may introduce neurocognitive processes beyond those sufficient for perception (Overgaard, 2004; Aru et al., 2012; Tsuchiya et al., 2015) and overestimate perception-related brain activity. Indeed, more recent paradigms capable of dissociating perception- and task-related brain activity (Panagiotaropoulos et al., 2012; Pitts et al., 2012; Shafto and Pitts, 2015; Kapoor et al., 2018; Cohen et al., 2020; Schlossmacher et al., 2021; Sergent et al., 2021) have consistently shown that under “no-report” conditions in which critical stimuli are perceived but not reported on a trial-by-trial basis, the P3b is abolished, but the PAN remains.

One such design uses inattentional blindness and independently manipulates perception and task-relevance of critical stimuli across different phases of the experiment (Pitts et al., 2012; Shafto and Pitts, 2015). In phase 1, participants remain naïve to critical stimuli and perform an orthogonal task. In phase 2, participants perform the same orthogonal task *after* being made aware of the critical stimuli. Finally, in phase 3, participants perform a task directly on the critical stimuli. Ideally, this design renders the critical stimuli perceived in phases 2 and 3 (but not phase 1), and task-relevant in phase 3 (but not phases 1 and 2). By comparing neural responses across phases 1 and 2, correlates of perception in the (near-)absence of task-relevance can be isolated. Similarly, by comparing responses across phases 2 and 3, correlates of task-relevance can be identified with few corresponding changes in perception. However, very few non-visual studies have utilized such a design (Schlossmacher et al., 2021).

Here, we adapted this design for use with sine-wave speech (SWS) (Remez et al., 1981; Remez, 2008), a synthetic audio signal comprising 3 to 4 frequency-modulated (FM) sine tones generated by removing natural speech cues (e.g. pitch) but preserving the time-varying frequencies and amplitudes of formants. Naïve SWS listeners typically report hearing “whistles”, “science-fiction” or other “computer-generated” sounds (Tuomainen et al., 2005; Davis and Johnsrude, 2007; Van Hedger et al., 2019; Cooke et al., 2022). However, listeners that know either the identity of the SWS utterance or, to a lesser extent, that the utterance is speech - often immediately recognize the speech content in SWS. Such a dramatic difference in perception of SWS despite zero changes to the stimulus lends itself well to the three-phase design discussed above. We recorded EEG during a three-phase experiment and predicted that a phonological version of the PAN would be elicited in phases 2 and 3, after participants were made aware of the identity of the SWS tokens, but not in phase 1, when participants remained naïve to the SWS. We further predicted that a P3b would be elicited by SWS tokens only in phase 3, when such tokens were task-relevant.

## Methods

### Ethics statement

All procedures were approved by the ethics committee at Reed College, and all participants gave written informed consent prior to their participation.

### Participants

30 undergraduate students of both sexes from Reed College between the ages of 18 and 23 participated in the study. All had normal or corrected-to-normal vision, no history of brain injury, self-reported normal hearing, and had no known history of neurological disorders. 12 were excluded from the study because they spontaneously became aware of the speech content in the SWS tokens during phase 1 (i.e. Noticers), and 1 was excluded because they failed to recognize the words in the SWS tokens after SWS training (i.e. Never noticed). This left 17 participants of both sexes in our final analysis (i.e. Non-noticers).

### Stimuli

The stimuli used for this study consisted of three pure tones (500 Hz, 1250 Hz, and 2000 Hz) and three SWS tokens (“brain”, “wave”, and “yard”) chosen based on pilot experiments as well as the fact that they avoid voiceless consonants that don’t translate well to SWS. The original words were recorded by a male speaker and converted to sine-wave speech using Praat (Boersma and van Heuven, 2001). Praat utilizes a formant tracker to detect formant frequencies and synthesize sine waves that track the center of these formants. Six additional SWS tokens (“chill”, “church”, “language”, “speech”, “world”, and “zombie”) served as “foil” stimuli during an intervening word recognition test between phases 1 and 2 of the main experiment (cf. Experimental Paradigm) but were not presented in any experimental phase. The pure tones were 600 ms in duration, while the SWS stimuli ranged from ∼480 to 600 ms in duration and all stimuli were presented at ∼46.5dB.

Control versions of the SWS tokens were constructed by “flipping” the frequencies of the second and third formants within a band- and time-limited region defined using amplitude thresholds specific to each token (Fig. 1). Specifically, for the lower frequency bound, we first identified the global maximum of the spectral magnitude, followed by identifying the adjacent local minimum. This minimum point served as our lower frequency bound for the spectral flipping procedure. The upper frequency bound was taken as the frequency point at which the spectral energy first dipped below 0.1% of the global maximum. Temporal bounds for the flipping procedure were defined in a similar manner, by thresholding the temporal envelope at 5% of the global maximum of the envelope signal (computed as the absolute value of the analytic signal, lowpass filtered below 100 Hz). After identifying these spectral and temporal bounds, the bounded signal was flipped using the time-domain algorithm proposed by Bell (Lyons, n.d.) and outlined in Lyons (Lyons, 2010) (Fig. 1A) and recombined with the remaining, unflipped portions of the signal. Importantly, this time-domain flipping method preserves phase information (which is not possible using methods based on the Fast-Fourier Transform and its inverse) and therefore does not introduce spectrotemporal discontinuities.

**Figure 1.**
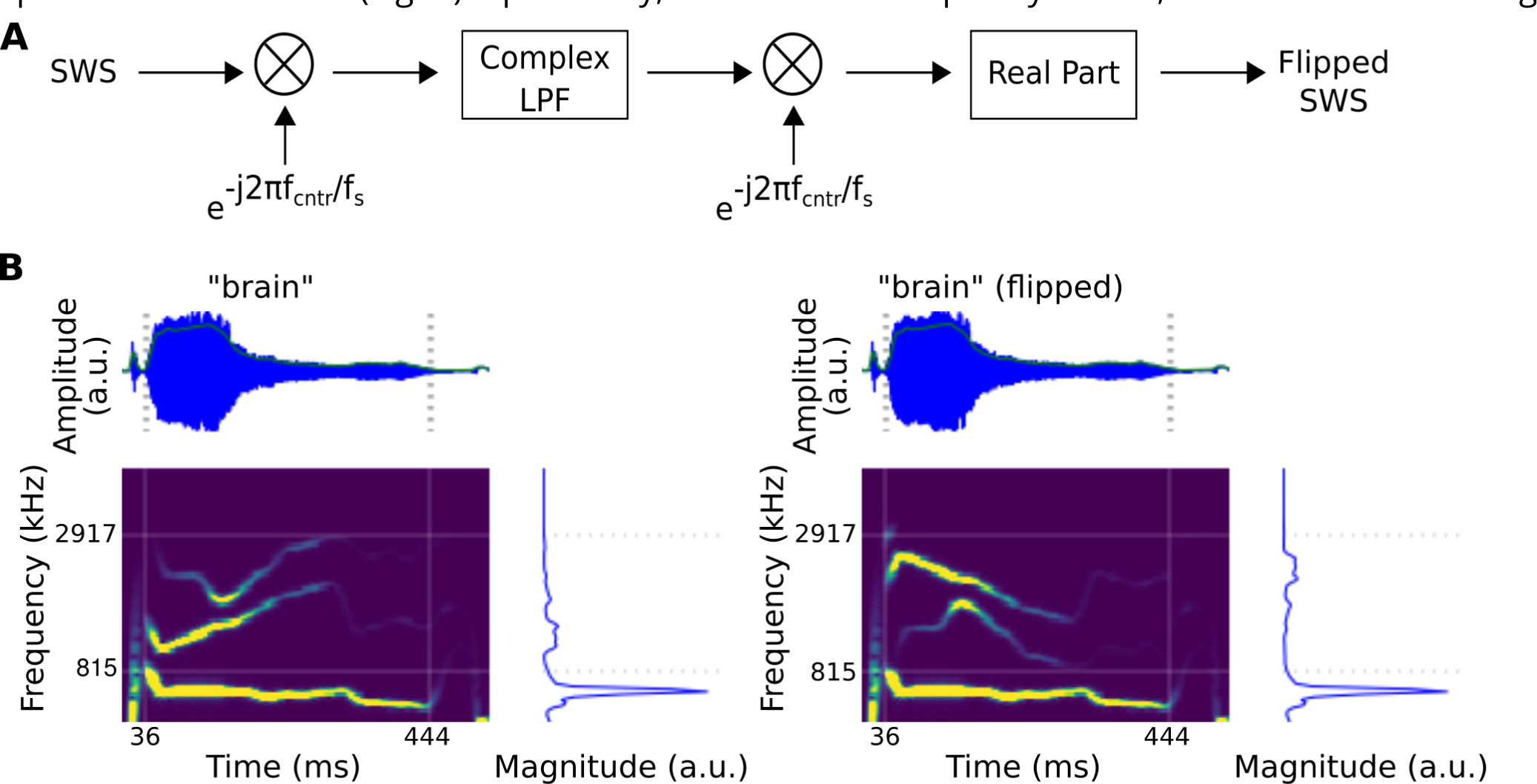
Flow diagram for creating frequency-flipped SWS tokens **(A)** and resulting SWS stimuli **(B). (A)** adapted from [39]. **(B)** Original SWS token for the word “brain” (left panels) and it’s flipped counterpart (right panels), where F2 and F3 have been flipped about a center frequency axis within a time- and band-limited region of the spectrogram. The resulting amplitude envelope (panels above spectrograms) and long-term spectra (panels to the right of spectrograms) are similar between original and flipped tokens.

This process resulted in “flipped” control SWS tokens that were otherwise identical to the normal SWS tokens (Fig. 1B) but not intelligible. Such a spectral manipulation is similar to spectral rotation first carried out by Blesser (Blesser, 1972) and used by several previous studies examining the neural basis of speech perception (Scott et al., 2000; Narain et al., 2003; Möttönen et al., 2006; Wild et al., 2012). However, our spectral manipulation is more subtle and in particular, preserves the spectral location and dynamics of the first, dominant formant (F1). Both the amplitude envelopes and long-term spectra are quite similar between flipped and unflipped SWS tokens. The main purpose of these control stimuli was to ensure that any neural differences observed between SWS stimuli in phase 2 (perceived as speech) versus phase 1 (perceived as noise) were not due to condition-order effects. In other words, because the control stimuli can only be perceived as noise, any neural changes between phase 1 and 2 for these stimuli can be ruled out as potential indices of conscious speech perception.

### Experimental paradigm

The experiment took place across three phases in which participants performed a category-selective 1-back memory task (on pure tones for phases 1 and 2 and on SWS tokens for phase 3). We presented each stimulus (3 SWS, 3 flipped controls, and 3 tones) 100 times in each phase (i.e., 300 SWS, 300 controls, 300 pure tones, 900 total stimuli per phase). 30% of the stimuli were one-back trials (90 stimuli of each category; 30 of each individual stimulus type), and these were discarded from EEG analysis. This left us with 210 trials per stimulus category, per phase, prior to artifact rejection. Across all three phases, hit rates, false-alarm rates, and reaction times to the one-back targets were measured. Presentation software (Neurobehavioral Systems, Berkeley, CA, USA) was used to control stimulus presentation and collect behavioral responses with an RB-830 Cedrus button box (Cedrus, San Pedro, CA, USA). To help subjects avoid eye movements, a fixation dot (1 deg visual angle) was presented constantly throughout the experimental trials.

In phase 1, participants performed a one-back task on pure tones (pressing a button whenever one of the three pure tones repeated on adjacent trials) and were unaware of the speech content within the SWS tokens. In phase 2, participants again performed the one-back task on the pure tones, but were no longer naïve to the speech content in the SWS tokens due to an intervening speech awareness assessment (cf. next paragraph). In phase 3, participants performed a one-back task on the SWS tokens, making them task-relevant. The control (frequency flipped) SWS stimuli were always task irrelevant.

Between phases 1 and 2, we assessed each listener’s perception of the stimuli using a speech awareness assessment. Specifically, participants were asked to provide confidence ratings regarding the extent to which they heard the following four categories of sounds: (i) distorted music, (ii) distorted words, (iii) distorted environmental sounds, or (iv) distorted animal sounds. Ratings ranged according to the following scale: (1) very confident I did not hear it, (2) confident I did not hear it, (3) uncertain, (4) confident I did hear it, and (5) very confident I did hear it. If any listener gave a confidence rating of 4 or 5 for hearing the “computer-generated noises” as “distorted words”, they were asked to write down any words they heard. Any participant who either gave a confidence rating of 4 or 5 or who identified any of the three words used as SWS in the study were excluded from further analysis (12 participants were deemed as Noticers, and excluded for this reason). After the speech awareness assessment, we then administered a training and recognition test. Participants were trained on 9 SWS tokens that included the three used in the study ( “brain”, “wave”, “yard”) as well as six additional SWS tokens that served as foils (“chill”, “church”, “language”, “speech”, “world”, and “zombie”). Participants listened to the SWS and original (i.e. natural) tokens in the order SWS -> Original -> SWS until the SWS tokens were clearly perceived as speech. A subsequent 10-way speech-recognition test (9 SWS words, 1 non-word) was administered to confirm that participants were able to recognize the SWS tokens.

The same speech awareness assessment that was administered after phase 1 was again administered after phase 2, but in this case, any participant who did *not* rate as a 4 or 5 their confidence in hearing “distorted words” were excluded from further analysis (1 participant, deemed as Never-noticed, was excluded for this reason).

### EEG recording

The EEG was recorded from 96 Ag/AgCl electrodes in an equidistant montage (EASYCAP, GmbH, Herrsching, Germany) using Brain Vision Analyzer (Brain Vision LLC, Morrisville, NC, USA) at a sampling rate of 500 Hz. Electrode impedances were kept below 5 kΩ. EEG data were filtered online with a 0.1 Hz highpass and a 150 Hz lowpass filter (both filters: 12 dB/oct roll-off) and amplified by three 32-channel amplifiers (Brain Amp Standard, Brain Products GmbH, Gilching, Germany). The online reference electrode was positioned at CPz. All recordings were conducted inside an electrically shielded, double-walled sound booth (Industrial Acoustics, Naperville, IL, USA) located in the Department of Psychology at Reed College. Eye movements and blinks were monitored by left and right horizontal EOG channels and a vertical EOG channel under the left eye, respectively. EEG data from all 30 participants were published on an OSF repository, including the 17 participants included in the analyses reported here, and the 13 participants excluded from analysis, along with behavioral data for all participants (https://osf.io/dsmjp/).

### Preprocessing

Preprocessing was done in Brain Vision Analyzer (Brain Vision LLC, Morrisville, NC, USA). EEG data were filtered offline with a 25 Hz low-pass filter (24 dB/oct roll-off), re-referenced to the common average reference, and epoched from -200 ms to 800 ms with respect to stimulus onset. Any epochs containing deflections larger than 70 µV were rejected. Overall, we retained 162 trials (on average) from 210 trials per phase and stimulus condition (22.8% rejection rate). All remaining epochs were baseline corrected using the mean signal in the pre-stimulus baseline period (-200 - 0 ms with respect to stimulus onset). All trials in which the same stimulus was repeated (i.e., one-back trials) were excluded from further analysis.

### Mass-univariate analysis

Mass-univariate analyses (MUA) (Maris and Oostenveld, 2007; Maris, 2012) were performed using the MNE-Python (Gramfort et al., 2014) using a combination of cluster-based two-way ANOVAs and, when necessary, cluster-based post-hoc paired comparisons. To examine effects of phase (phase 1, phase 2, phase 3) and stimulus type (SWS, flipped SWS), we carried out two, two-way ANOVAs: (i) Phase 1 versus Phase 2 for SWS and flipped SWS, (ii) Phase 2 versus Phase 3 for SWS and flipped SWS. In light of the significant clusters found for the interaction component of the first ANOVA (phases 1 and 2, SWS and flipped SWS; cf. Results), subsequent paired comparisons were made (i) between phases (phase 1 versus phase 2) for SWS, (ii) between phases (phase 1 versus phase 2) for flipped SWS, (iii) between stimulation types (SWS versus flipped SWS) in phase 1, and (iv) between stimulation types (SWS versus flipped SWS) in phase 2. Our first hypothesis was that there would be significant differences in early latency ranges between phase 2 and phase 1 for SWS, but not for “flipped” SWS, reflecting the difference in how SWS tokens, but not “flipped” SWS tokens, are perceived in phase 2 (as speech) versus phase 1 (as noise). Our second hypothesis was that a P3b would only be observed in phase 3, and would therefore appear as a difference in later latency ranges between phase 3 and phase 2 for SWS, but not “flipped” SWS. For each of the comparisons, correction for multiple comparisons was carried out using the cluster-based permutation. The cluster-level *F*-statistic or *T*-statistic was taken as the sum of contiguous test-statistics (i.e. *F*- or *T*-values) when sample-level *p*-values were less than 0.05. A continuous spatial-temporal cluster was defined by adjacent channels at continuous temporal intervals. Channel adjacency was calculated using Delaunay triangulation based on 2D channel locations (Smith and Nichols, 2009). To determine significance of the cluster-level statistic, condition labels were shuffled 10,000 times to produce a null distribution of cluster-level statistics. Cluster summary statistics were compared to the null distribution, and clusters for which summary statistics exceeded the 95th percentile of the null distribution were considered significant (corresponding to *p* < 0.05 at the cluster level) (Maris and Oostenveld, 2007; Maris, 2012).

### Bayes factors

In contrast to the frequentist approach of null-hypothesis significance testing (NHST) where non-significant tests cannot be used as evidence in favor of the null hypothesis, Bayes factors can be used to interpret data as being evidence for or against the null hypothesis (Dienes, 2014; Dienes and Mclatchie, 2018; Wagenmakers et al., 2018). Here, we computed Bayes factors on behavioral data as well as clustered, windowed, and averaged ERP responses using paired versions of the ttestBF function in the BayesFactor package available for R (Morey, 2023). The default assumptions for this function include that the true standardized difference is 0 under the null hypothesis and Cauchy-distributed with a scale = r = √2/2 under the alternative hypothesis.

### Multivariate pattern analysis

MVPA was performed in MNE-Python using ERP data across all 96 channels, filtered between 0.05 and 20 Hz and resampled at 50 Hz. Stimulus type (SWS versus flipped SWS) was decoded within each of the three phases (phase 1, phase 2, phase 3) separately. Our hypothesis was that classification of stimulus type in phase 1 should be at chance levels due to the fact that all 17 of the participants included in our analysis reported hearing these stimuli as noise in this initial phase of the experiment. In contrast, after participants were made aware of the SWS word tokens and thus perceived them as speech, classification between SWS and flipped SWS in phase 2 should be significantly above chance, at least during some portion of the time window. Furthermore, when the SWS tokens were made task-relevant in phase 3, we expected further enhancements of classification accuracy between SWS and flipped SWS. Classification differences of SWS versus flipped SWS tokens between phases 1 and 2 should reveal information in the ERPs about perceptual differences; classification differences between phases 2 and 3 should reveal information in the ERPs about task-relevance. We used support vector machines (SVM) as the decoding model (linear kernel function, C=1), and input the amplitude values over all 96 channels of the respective ERPs at each time point to the models. We used 3-fold cross validation and reported the average area under the curve (AUC) of the receiver operating characteristic across the three folds (chance level = 0.5) at each time point. Cluster-based permutation tests were used to correct for multiple comparisons and test for temporal windows of above-chance decoding AUC (p < 0.05 at the cluster level) (Maris and Oostenveld, 2007; Maris, 2012; Bae and Luck, 2019). The sample-level *T*-statistics were derived from one sample *t*-tests for above-chance mean decoding AUC across participants at each time point. The cluster-level *T*-statistic was taken as the sum of contiguous *T*-values when sample-level *p*-values were less than 0.05. To determine significance of the cluster-level *T*-statistic, condition labels were shuffled 10,000 times to produce a null distribution of cluster-level *T*-statistics. Cluster summary statistics were compared to the null distribution, and clusters whose cluster-level statistic exceeded the 95th percentile of the null distribution were considered significant (corresponding to a cluster-level *p*-value < 0.05). Similarly, we performed a 3-fold temporal generalization (TG) analysis (King and Dehaene, 2014; King et al., 2014) across all three phases between SWS versus flipped SWS. 10,000 times cluster-based permutation were also used to correct for multiple comparisons for nearby time points on temporal generalized maps. We also report the average multivariate weights map across all 3-fold models for each phase decoding (Haufe et al., 2014).

## Results

### Behavior

In phases 1 and 2, participants performed a 1-back detection task on pure tones in order to (i) maintain their attention to stimuli and (ii) render SWS tokens task-irrelevant. On average, participants detected 99% of such pure-tone repetitions in phase 1 (mean ± 95% confidence interval = 99.15 ± 0.77) and 97% of such pure-tone repetitions in phase 2 (97.06 ± 1.47). This difference, though small, was statistically significant (*T*_16_ = 2.74, p < 0.05) (BF_10_ = 3.98). False alarms were more common in phase 2 (0.37 ± 0.34%) than phase 1 (0.04 ± 0.03%) but remained low overall and did not differ significantly between the two phases (*T*_16_ = -2.0, n.s.), although weak evidence was found in favor of a difference in false-alarm rates between phases (BF_10_ = 1.24). Reaction times did not significantly differ between phase 1 (277.49 ± 36.56 ms) and phase 2 (284.73 ± 44.22 ms) (*T*_16_ = -0.73, n.s.) (BF_10_ = 0.32).

After each of the first two phases, participants were asked to report their confidence in hearing distorted music, distorted words, distorted environmental sounds, or distorted animal sounds (Fig. 2). Across all participants in our final analysis (N=17), none gave a confidence rating greater than 3 for hearing distorted words in phase 1. Confidence ratings for other categories varied. Similarly, only one of the 17 participants reported hearing words in phase 1, and none of the words identified by that participant included any of the three SWS tokens that were present in the stimuli. Thus, we can be confident that none of the 17 participants in our final analysis heard the words in the SWS stimuli in phase 1.

**Figure 2.**
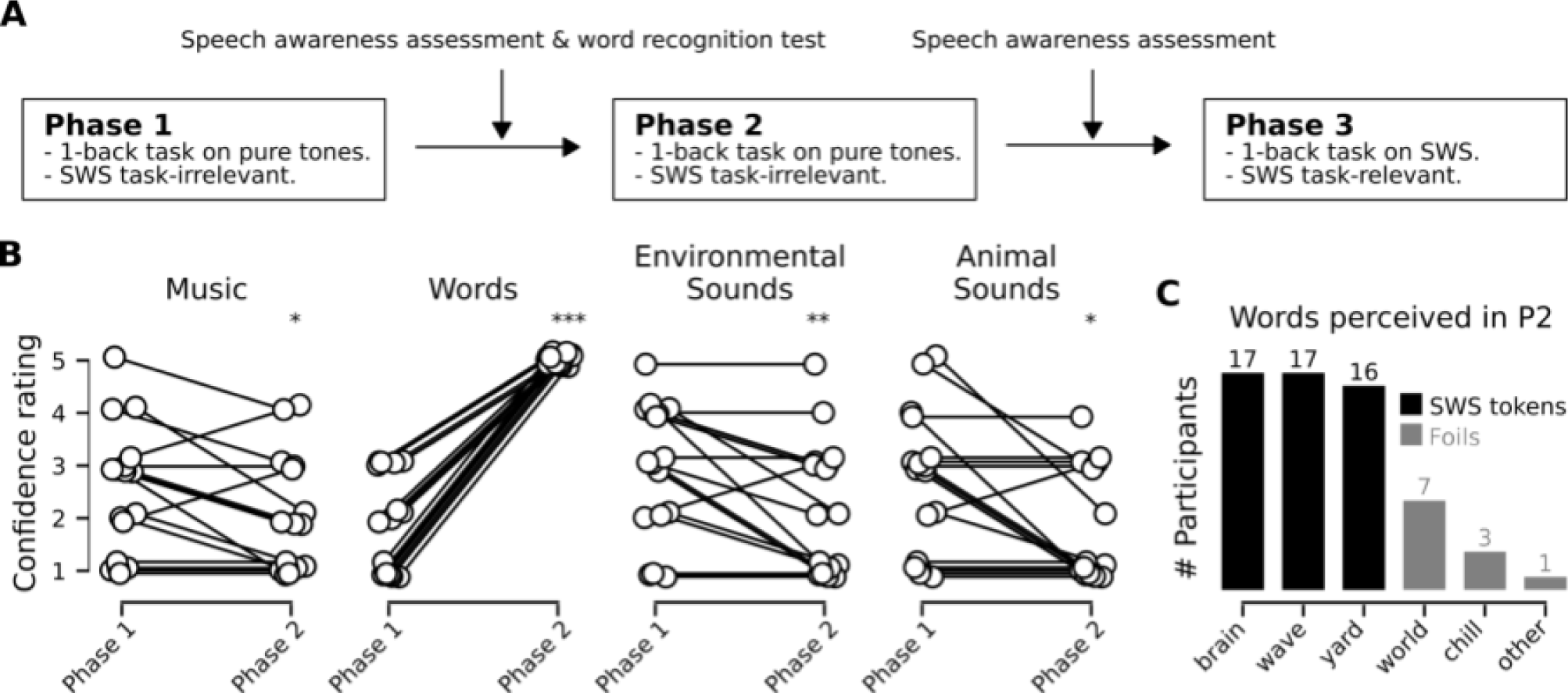
Three-phase sine-wave speech paradigm (A) and behavioral results (B, C). (A) SWS tokens were task-irrelevant in phases 1 and 2, when participants performed a 1 -back task on pure tones. All 17 participants included were naive to the presence of speech in phase 1, but informed in phase 2 due to an intervening speech awareness assessment and training on SWS. The same awareness assessment was given after phase 2 to confirm participants heard speech in that phase before performing a 1-back task on the SWS tokens in phase 3, making them task relevant. (B) Rating scores across phases 1 and 2 for four different sound categories. Only words showed a positive-going significant difference between phases 1 and 2, confirming the effectiveness of the intervening awareness assessment and training on SWS. All other categories showed significantly decreased rating scores between phases 1 and 2. * p < 0.05; ** p < 0.01; *** p < 0.001 (C) Number of participants who included specific words in response to free-form questioning after phase 2.

After phase 1, we also administered a SWS training session and a speech recognition test to ensure that participants were able to identify the SWS tokens (cf. Methods). After the training session, participants correctly identified 94% of all nine SWS tokens, and 99% of the three SWS tokens (“brain”, “wave”, “yard”) that served as the critical stimuli in phases 1, 2, and 3 of the main EEG experiment. Due to this training and recognition testing after phase 1, the speech awareness assessment administered after phase 2 resulted in confidence ratings for hearing distorted words at ceiling-levels for all 17 participants included in our final analysis (Fig. 2), and all but one listener reported hearing the specific tokens “brain”, “wave”, and “yard” during phase 2 (the remaining listener heard both “brain” and “wave” but not “yard”). The mean difference in confidence ratings for words was 3.18 ± 0.45 (mean ± 95% confidence interval). Thus, while participants most likely did not hear any words in phase 1, they almost certainly heard the three specific words in phase 2 (*T*_16_ = 14.84, *p* << 0.001) (BF_10_ >> 100), even though the stimuli were physically identical across phases. Confidence ratings for other categories remained variable but did decrease significantly between phases 1 and 2 for music (*T*_15_ = -2.52, *p* < 0.05) (BF_10_ = 2.73), environmental sounds (*T*_15_ = -3.15, *p* < 0.01) (BF_10_ = 7.70), and animal sounds (*T*_15_ = -2.78, *p* < 0.05) (BF_10_ = 4.16). The corresponding mean differences in confidence ratings were -0.56 ± 0.46 (music), -0.69 ± 0.45 (environmental sounds), and -0.88 ± 0.65 (animal sounds).

In phase 3, the participants performed a 1-back task on the SWS tokens, making them task-relevant. Listeners detected nearly 94% of such 1-back SWS tokens (93.55 ± 3.32) (mean ± 95% confidence interval), but hit rates in phase 3 were not as high as the hit rate for tones in phase 1 (*T*_16_ = -3.95, *p* < 0.01) (BF_10_ = 33.59) or phase 2 (*T*_16_ = -2.61, *p* < 0.05) (BF_10_ = 3.17). False alarm rates were significantly higher for the 1-back task in phase 3 (2.97 ± 2.23) as compared to phase 1 (*T*_16_ = 2.78, *p* < 0.05) (BF_10_ = 4.22) or phase 2 (*T*_16_ = 2.60, *p* < 0.05) (BF_10_ = 3.11), but remained low overall. We attribute the higher false-alarm rate in phase 3 to the fact that the “flipped” control versions of the SWS tokens were perceptually more similar to the non-flipped tokens than the pure tones were to either the flipped or unflipped SWS tokens. In other words, the 1-back task was easier in phase 1 and 2 than in phase 3.

### EEG

Because our hypothesis was that we should only observe differences in ERPs between phases for SWS tokens (and not frequency-flipped SWS tokens), and because we didn’t have strong a-*priori* hypotheses about when or where such differences would manifest, we first carried out a cluster-based two-way ANOVA for each contrast of interest, each with two factors, where each factor had two levels. Both ANOVAs included factors of phase (either phase 2 versus phase 1 or phase 3 versus phase 2) and stimulation type (SWS versus flipped SWS). The first ANOVA (phase 2 versus phase 1 and SWS versus flipped SWS) revealed a single significant cluster for the two-way interaction (Fig. 3A).

**Figure 3.**
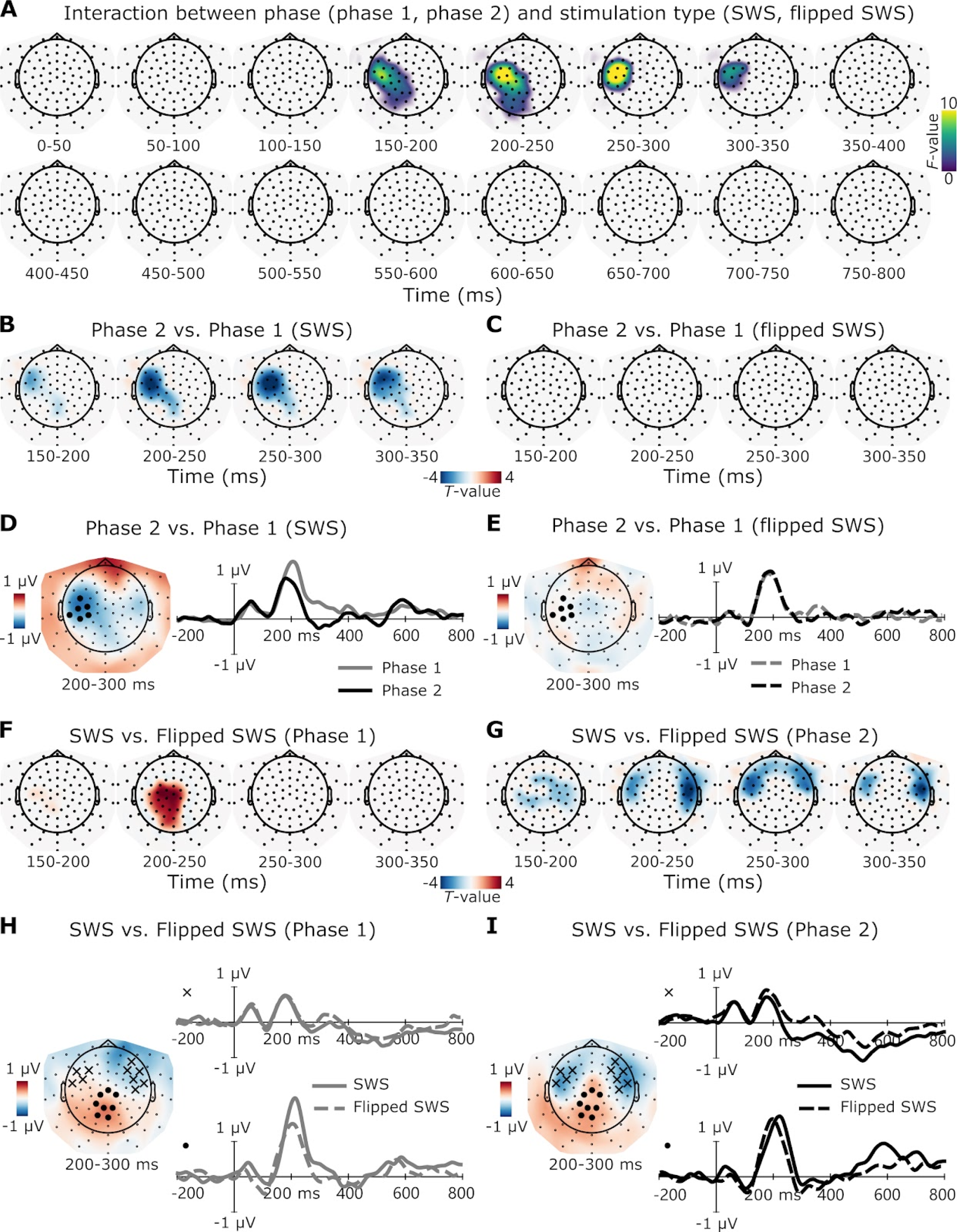
Event-related potentials and mass-univariate analyses for SWS and flipped SWS for phases 1 and 2. Significant clusters are plotted with their corresponding *F-* or 7-values; non-significant areas are left blank. For all the scalp topographies, waveforms are derived from electrode clusters represented by black dots or the symbol ’x’. For all waveform plots, solid traces represent ERPs in response to SWS tokens, and dashed traces represent ERPs in response to flipped SWS tokens. **(A)** A significant interaction between phase (phase 1 vs. 2) and stimulus type (SWS vs. flipped SWS) was observed between -150 and 350 ms. (B, C) Significant clusters derived from paired t-tests across phases in SWS and flipped SWS tokens. (B) Only the SWS tokens showed significant differences in the ERPs across phases, between -150 and 350 ms. (C) Any differences observed for flipped SWS were not statistically significant. **(D, E)** Scalp topographies (Phase 2 - Phase 1) and waveforms for SWS tokens **(D)** and flipped SWS tokens **(E). (F, G)** Significant clusters derived from paired t-tests across stimulation types in phases 1 and 2. **(F)** In phase 1 a significant different ERPs between SWS and flipped SWS were observed, between -200 and 250 ms over posterocentral scalp sites. (G) In phase 2 a significant difference between SWS and flipped SWS was observed between -150 and 350 ms over anterolateral scalp sites. **(H, I)** Scalp topographies (SWS - flipped SWS) and waveforms for phase 1 **(H)** and phase 2 **(I).** Symbols in (H) and (I) indicate from which scalp sites the waveforms are from.

Subsequent cluster-based paired *t*-tests revealed a single significant cluster across phases for SWS tokens (Fig. 3B), which were perceived as speech in phase 2 but as noise in phase 1. No such cluster was found for flipped SWS tokens, which were perceived as noise in both phases (Fig. 3C). This difference between phase 2 and phase 1 for unflipped SWS was clear in both the ERP topography and waveforms as a difference between ∼200-300 ms over left anterocentral scalp sites (Fig. 3D). The size of this effect, as measured by the difference (phase 2 - phase 1) in amplitude of the clustered waveforms between 150 and 350 ms, was -0.54 ± 0.26 µV (mean ± 95% confidence interval) (Cumming, 2014). By contrast, no such difference was observed for the frequency-flipped SWS tokens (Fig. 3E) (effect size: -0.06 ± 0.21). To determine whether the evidence in each case supports either the null hypothesis (no difference between phases) or the alternative hypothesis, we used Bayes factors. The average difference in amplitude between phase 2 and phase 1 from 150 - 350 ms for SWS (Fig. 3D) yielded a Bayes factor of BF_10_ = 71.26, indicating strong evidence in favor of a difference between the two phases for SWS. In contrast, the average difference in amplitude between phase 2 and phase 1 from 150 - 350 ms for flipped SWS (Fig. 3E) yielded a Bayes factor of BF_10_ = 0.30, indicating substantial evidence in favor of no difference between the two phases for flipped SWS.

Cluster-based paired *t*-tests between SWS and flipped SWS *within* phases showed two significant clusters from different areas depending on phase. In phase 1 (Fig. 3F), a brief difference was observed from ∼190 to 250 ms over posterocentral scalp sites (a difference topography between SWS and flipped SWS and corresponding waveforms for phase 1 is shown in Fig. 3H). Note here that this posterocentral difference between SWS and flipped SWS in phase 1, while not significantly different in phase 2, showed a similar numerical difference in scalp topographies across the phases (i.e., compare topographies in Figs. 3H and 3I). In phase 2, a longer-lasting difference over lateralized anterocentral scalp sites was observed from approximately 150 to 350 ms (Fig. 3G) (a difference topography between SWS and flipped SWS and corresponding waveforms for phase 2 is shown in Fig. 3I). Note that while this difference was bilateral, the left-lateralized location of the difference between SWS and flipped SWS tokens in phase 2 was similar to the difference between phases 1 and 2 for SWS (compare Fig. 3B and Fig. 3G as well as Fig. 3D and Fig. 3I). The size of these effects, as measured by the difference (SWS - flipped SWS) in amplitude of the clustered waveforms between 150 and 350 ms, was -0.14 ± 0.20 (mean ± 95% confidence interval) in phase 1 (Fig. 3H, top traces), and -0.45 ± 0.21 in phase 2 (Fig. 3I, top traces). The corresponding Bayes factors were BF_10_ = 0.58 in phase 1 and BF_10_ = 82.08 in phase 2, indicating weak evidence for no difference in phase 1 and strong evidence for a difference in phase 2.

For phase 3 vs phase 2 (SWS perceived as speech in both phases, but task-*relevant* in phase 3 and task-*irrelevant* in phase 2), no significant clusters were observed for the two-way interaction between phase (phase 3 versus phase 2) and stimulation type (SWS versus flipped SWS) (data not shown). However, we did observe significant clusters for the main effects of phase (phase 3 versus phase 2) and stimulation type (SWS versus flipped SWS).

For phase (Fig. 4A), three significant clusters were observed: (i) a phase 3 > phase 2 posterocentral (near Pz, but slightly left-lateralized) positivity between ∼400 and 800 ms, (ii) a phase 3 > phase 2 anterocentral (near FCz) negativity between ∼400 and 800 ms, and (iii) an early difference over central (near Cz) scalp sites between ∼0 and 350 ms. For the later clusters, these effects were numerically larger for SWS tokens than for flipped SWS tokens (Figs. 4C and 4D), possibly reflecting the task-relevance (in Phase 3) of the SWS tokens but not flipped SWS tokens. However, as stated above, no significant interaction was observed between phase and stimulation type (cf. Discussion).

**Figure 4.**
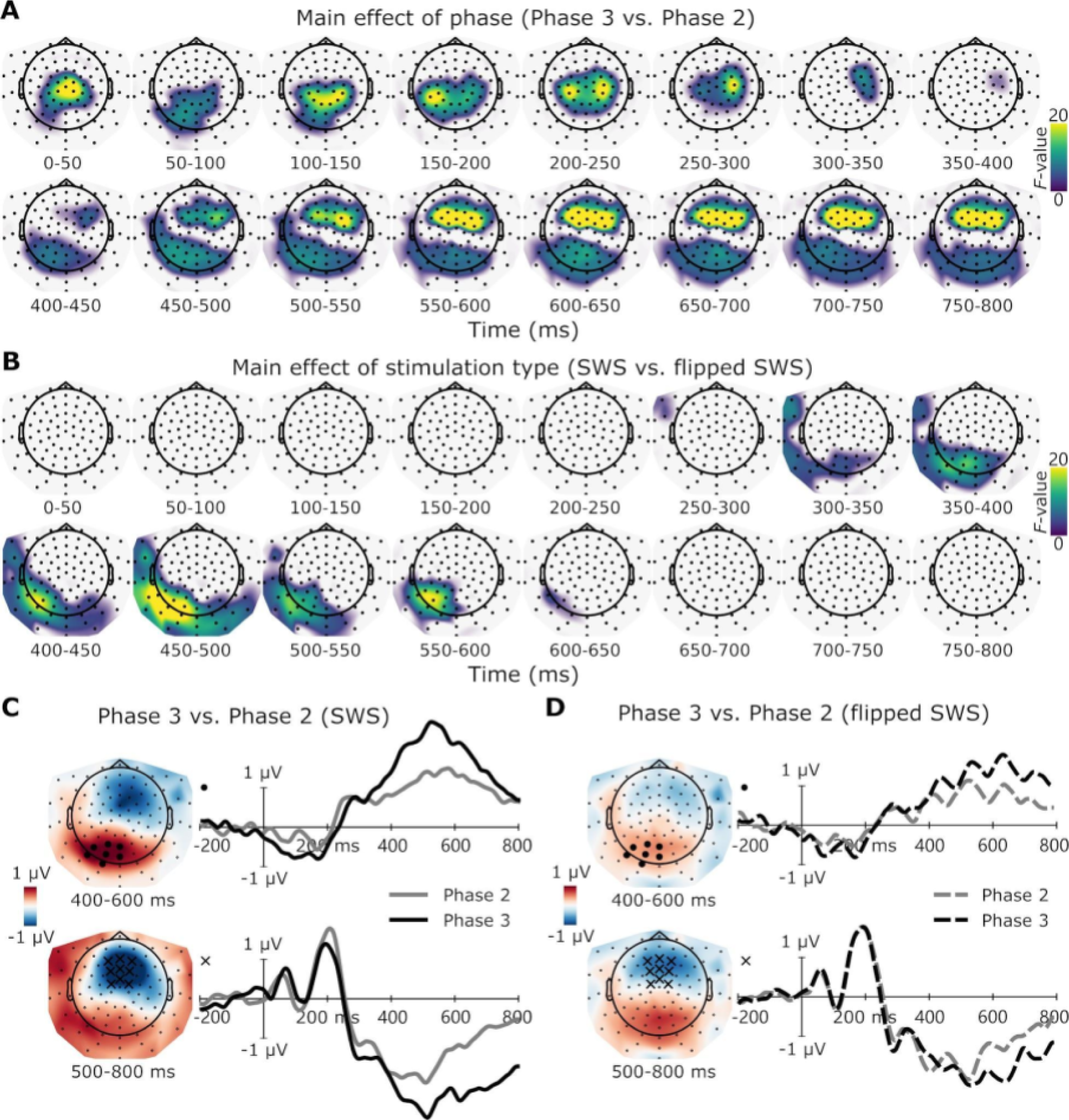
Results from a mass-univariate two-way ANOVA **(A** and **B)** and event-related potentials **(C** and **D)** for SWS and flipped SWS and phases 2 and 3. **(A)** Significant main effects of phase across phases 2 and 3 was observed from 0 to 350 ms and from -400 to 800 ms. (B) Significant main effect of stimulation type between SWS and flipped SWS was observed from -350 to 600 ms. **(C, D)** Scalp topographies (Phase 3 - Phase 2) and waveforms for SWS tokens (C) and flipped SWS tokens (D). Symbols in (C) and (D) indicate which scalp sites the waveforms are from.

For stimulation type (Fig. 4B), SWS stimuli generally showed larger responses in both phases than did flipped SWS stimuli (compare black and gray traces across panels C and D in Fig. 4). This effect appeared to be numerically larger for phase 3 than phase 2, but again, no significant two-way interaction between phase and stimulation type was observed. Finally, several earlier task-related effects were also evident in the Phase 3 versus 2 contrast for SWS stimuli, possibly reflecting task-based differences.

### Multivariate pattern analysis

To further examine potential differences between conditions, we performed multivariate pattern analysis (MVPA) on single trials (Fig. 5). We used temporal generalization (Fig. 5A) and temporal decoding (i.e., the diagonal of the temporal generalization matrices) (Fig. 5B) to carry out three different classification analyses (SWS versus flipped SWS stimuli within phase 1, phase 2 and phase 3). If there are any significant differences between SWS and flipped SWS, classification AUC should be significantly higher than chance at those time points (temporal decoding along the diagonal) or matrix entries (temporal generalization). For both the diagonal and temporal generalization, we hypothesized a larger extent of above-chance decoding for phase 2 versus phase 1 and for phase 3 versus phase 2.

No significant decoding between SWS and flipped SWS was observed in phase 1 (Fig. 5A & 5B, left panels). In phase 2, AUC was significantly higher than chance between approximately 220 and 580 ms on the diagonal, with maximal decodability around 300 ms (Fig. 5B, middle panel). Temporal generalization of this pattern was mostly confined to entries around the diagonal (Fig. 5A, middle panel), and any patterns that did generalize off-diagonal entries were relatively early compared to the pattern observed in phase 3. In phase 3, AUC was significantly higher than chance from approximately 160 ms to the end of the epoch (Fig. 5B, right panel). Furthermore, in contrast to phase 2, generalization of the patterns decoded in phase 3 extended far off the diagonal, particularly for later training/testing times (Fig. 5A, right panel).

**Figure 5.**
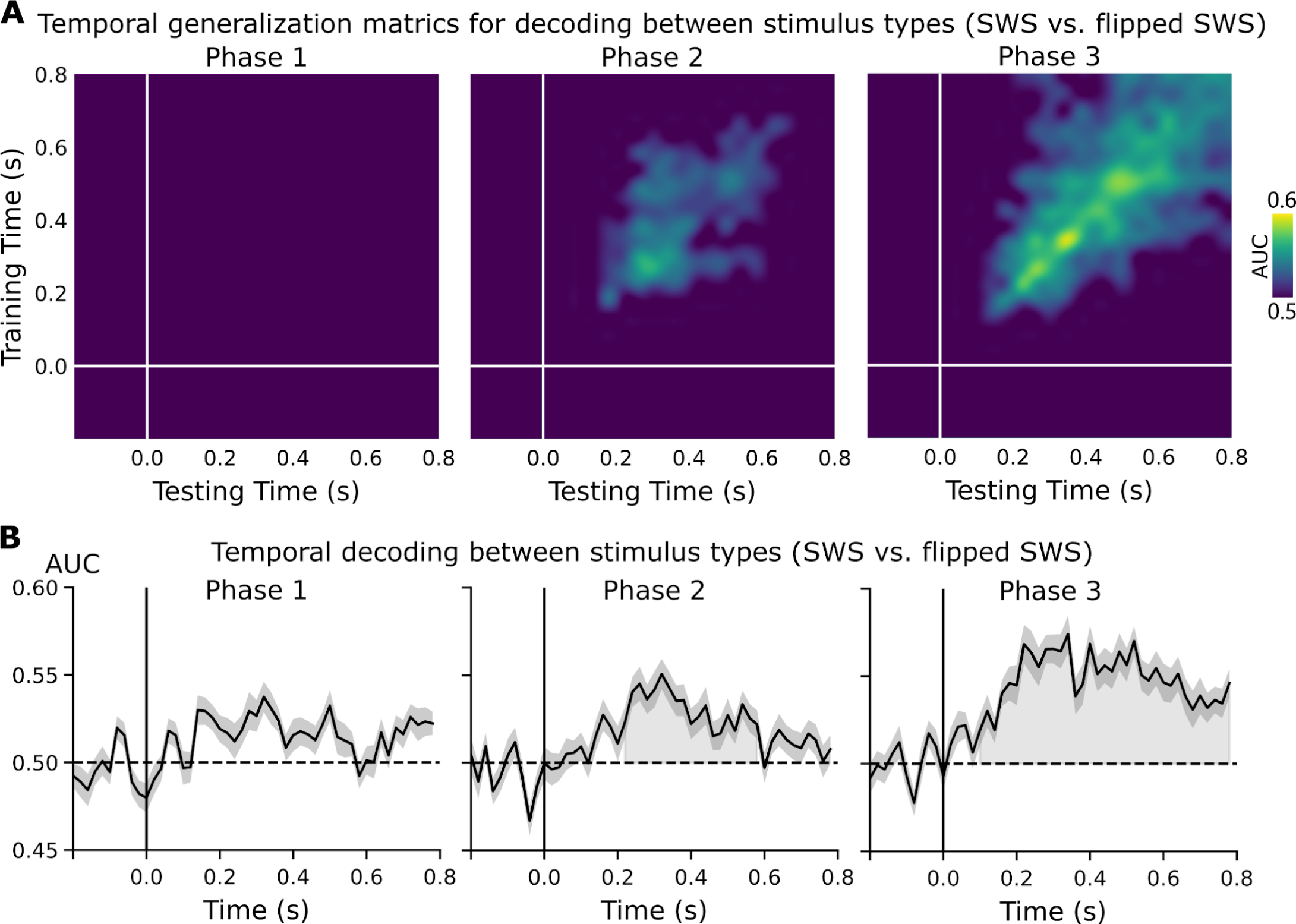
Decoding AUC of SWS versus flipped SWS in phases 1, 2, 3. **(A)** All non-significant matrix entries are reported as AUC values equivalent to the chance level (purple), and only significant pixels are reported with their corresponding AUC values. (B) The black trace represents the averaged AUC across participants, and deep gray shade represents the AUC standard error across participants. Significant clusters were plotted with light gray shading under the AUC trace. decoding (i.e., the diagonal of the temporal generalization matrices) (Fig. 5B) to carry out three different classification analyses (SWS versus flipped SWS stimuli within phase 1, phase 2 and phase 3). If there are any significant differences between SWS and flipped SWS, classification AUC should be significantly higher than chance at those time points (temporal decoding along the diagonal) or matrix entries (temporal generalization). For both the diagonal and temporal generalization, we hypothesized a larger extent of above-chance decoding for phase 2 versus phase 1 and for phase 3 versus phase 2.

**Fig. 6.**
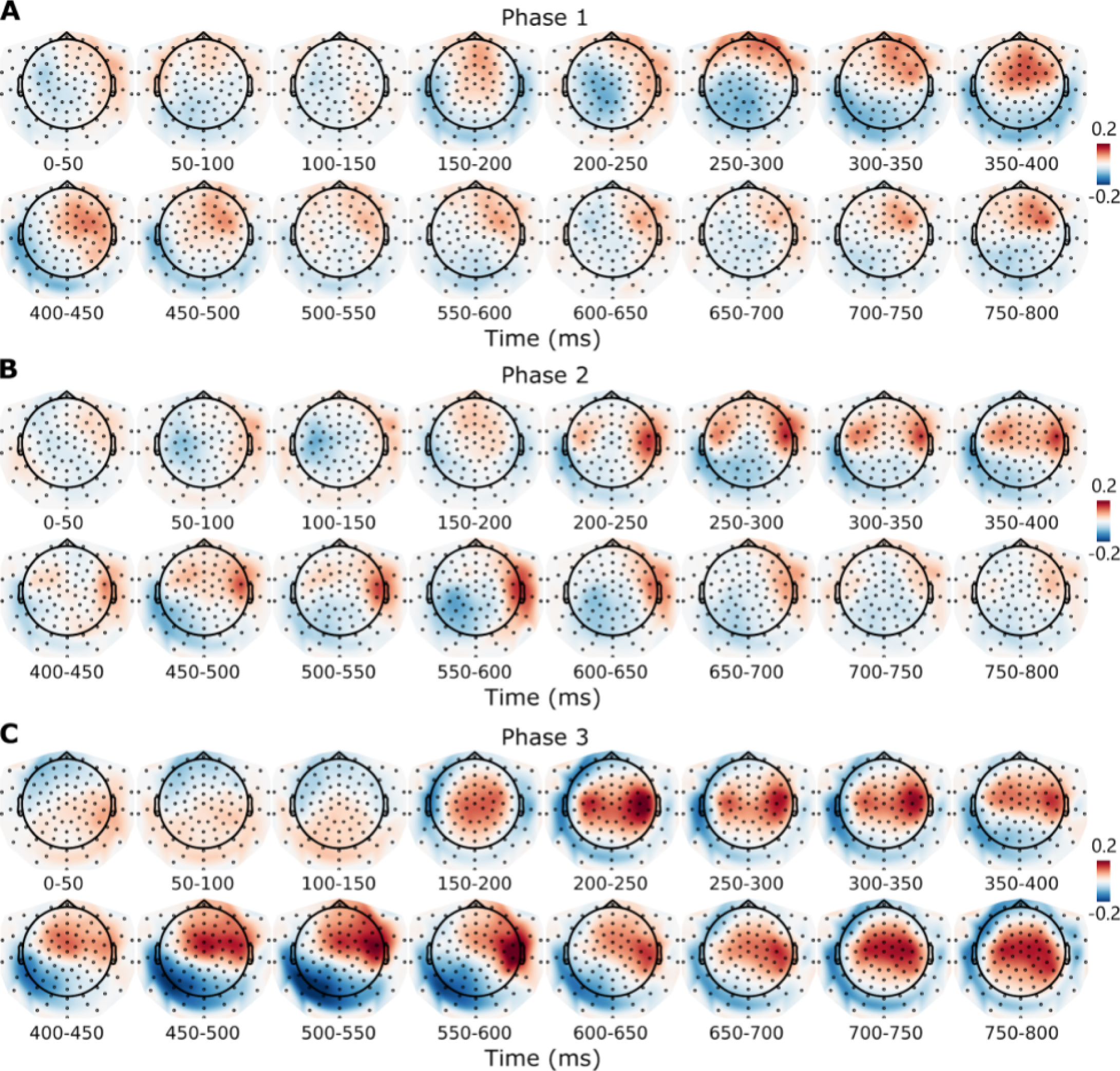
shows the averaged MVPA feature weights of all three folds across all participants of SWS versus flipped SWS in phases 1, 2, and 3. Feature weight patterns were relatively weak in phase 1 (Fig. 6A), including some anterior scalp sites around 300 ms. In phase 2 (Fig. 6B) a strong pattern was observed bilaterally at anterolateral scalp sites from approximately 200 to 400 ms. This pattern is similar to the *T-* statistic map between SWS and flipped SWS in phase 2 (Fig. 3G). Here, however, the difference persisted longer (until ∼600 ms) in right temporal electrodes until 600ms. In phase 3, an even stronger pattern than phase 2 was observed bilaterally from approximately 200 to 400 ms, with progression to right temporal electrodes until 600ms and to central electrodes until the end of the epoch (Fig. 6C).

No significant decoding between SWS and flipped SWS was observed in phase 1 (Fig. 5A & 5B, left panels). In phase 2, AUC was significantly higher than chance between approximately 220 and 580 ms on the diagonal, with maximal decodability around 300 ms (Fig. 5B, middle panel). Temporal generalization of this pattern was mostly confined to entries around the diagonal (Fig. 5A, middle panel), and any patterns that did generalize off-diagonal entries were relatively early compared to the pattern observed in phase 3. In phase 3, AUC was significantly higher than chance from approximately 160 ms to the end of the epoch (Fig. 5B, right panel). Furthermore, in contrast to phase 2, generalization of the patterns decoded in phase 3 extended far off the diagonal, particularly for later training/testing times (Fig. 5A, right panel).

## Discussion

We recorded ERPs to SWS words in a three-phase paradigm in which we independently manipulated perception and task-relevance. In phase 1, listeners were naive to the presence of speech and heard SWS as noise or whistles. In phase 2, after an intervening awareness assessment and SWS training, listeners were aware of the presence of speech and heard SWS as speech. In both phases, SWS was task-irrelevant and listeners performed an orthogonal 1-back task on pure tones. In phase 3, after another awareness assessment, listeners again heard SWS as speech but performed a 1-back task on the now-task-relevant words. We observed a novel phonological version of a perceptual awareness negativity and replicated earlier findings showing that the P3b is associated more with task performance than perceptual awareness *per se*.

### Perceptual awareness negativity

Perception-related activity for SWS between phase 2 (task-irrelevant, perceived as speech) and phase 1 (task-irrelevant, not perceived as speech) as well as between SWS and flipped SWS in phase 2 was observed as (i) frontocentral scalp negativities from ∼200 – 300 ms (Fig. 3) and (ii) significantly above-chance decoding between SWS and flipped SWS in phase 2 from ∼200 – 600 ms (Fig. 5). No such differences were observed when comparing either flipped SWS (always perceived as noise) between phases 1 and 2 or when comparing SWS and flipped SWS within phase 1 (both perceived as noise).

The early ERP differences we observed between SWS in phases 1 and 2 as well as between SWS and flipped SWS in phase 2 can be interpreted as a phonological version of a perceptual awareness negativity (Dembski et al., 2021), a negative-going response in modality-specific sensory cortex that co-varies with perceptual awareness. Prior work in audition has identified such correlates for simple stimuli (i.e. tones) (Gutschalk et al., 2008; Eklund and Wiens, 2019; Fernandez Pujol et al., 2023), but few have used designs that can dissociate perception from task relevance (Hillyard et al., 1971; Wiegand et al., 2018; Sergent et al., 2021). Studies that have employed no-report paradigms in audition have found an AAN when participants were aware of (and could identify) critical word stimuli (Schlossmacher et al., 2021). The scalp topography of the AAN reported in that study was less lateralized than the one observed here. This could reflect idiosyncratic samples or a bilateral effect in our case that was simply stronger over left versus right scalp sites for the contrast between phases 1 and 2 (although the effect was bilateral for the contrast between SWS and flipped SWS within phase 2).

The negativity observed here is consistent with the idea that the contents of conscious perception arise from specialized processing in modality-specific cortex (Koch et al., 2016; Boly et al., 2017; Lamme, 2020; Dembski et al., 2021) [but see (Bola and Doradzińska, 2021; Doradzińska and Bola, 2023) for an alternative, attention-based view]. Future work could seek to rule out possible contributions of attention or word identification (Koivisto et al., 2017) and identify the generators of these effects to determine how consistent they are with prior neuroanatomical studies of speech perception (Hickok and Poeppel, 2015; Scott, 2019).

### Neural correlates of speech perception

Prior studies using SWS have observed perception-related differences at several levels of auditory processing, including auditory cortex as measured by functional MRI (Dehaene-Lambertz et al., 2005; Möttönen et al., 2006), high gamma-band activity in speech-motor cortex as measured by electrocorticography (Khoshkhoo et al., 2018), and even subcortical processing as measured by the frequency following response (FFR) (Cheng et al., 2021). However, as opposed to the contrasts made here, which were controlled for order and task-relevance, prior studies using SWS have contrasted (i) task-relevant SWS perceived as speech versus task-irrelevant SWS perceived as noise (Khoshkhoo et al., 2018), (ii) task-relevant SWS perceived as speech versus task-relevant SWS perceived as noise (Dehaene-Lambertz et al., 2005; Möttönen et al., 2006), or (iii) task-irrelevant SWS perceived as noise versus task-irrelevant SWS perceived as speech, without within-subject control stimuli (Cheng et al., 2021). These methodological details are important to consider.

For example, one study recorded electrocorticography in response to SWS sentences both before and after exposure to corresponding natural speech (Khoshkhoo et al., 2018). Compared to pre-training, SWS sentences post-training elicited larger high gamma-band activity over speech-motor areas of frontal cortex. However, patients were required to identify particular words in the sentences after but not before training, making It unclear whether this activity was associated with differences in perception or the task. Few changes were observed in either superior temporal gyrus or superior temporal sulcus, regions thought to be involved in perception of (sine-wave) speech from functional MRI studies (Dehaene-Lambertz et al., 2005; Möttönen et al., 2006). Adapting our three-phase design for use with electrocorticography (or other more spatially resolved methods) could resolve these discrepancies and help identify the generators of the PAN reported here.

Another study recorded frequency-following responses (FFR) to SWS across two sessions in two groups of participants during passive listening (Cheng et al., 2021). One group received SWS training between sessions; the other did not. Compared to the control group, the group who received SWS training showed more robust formant tracking after versus before training. This suggests that even *sub*-cortical representation of SWS stimuli is enhanced via perception of the tokens as speech, particularly given that the frequency of the formants being tracked in that study (> 300 Hz) were well above the typical limit of FFRs in cortex (∼100 Hz) (Tichko and Skoe, 2017; Bidelman, 2018) [but see (Hamilton et al., 2021)]. Such enhancement of subcortical responses could occur through corticofugal projections (Winer, 2005; Asilador and Llano, 2021; Lesicko and Geffen, 2022). However, without incorporating control stimuli into the design (e.g., flipped SWS), it is difficult to determine whether the neural changes observed by Cheng et al. (2021) were due to changes in perception or to something more generic such as changes in expectations or attention or differential exposure to the stimuli during training.

Finally, other studies using noise-vocoded speech (Shannon et al., 1995) - a stimulus that, like SWS, is initially difficult to hear as speech (Davis et al., 2005; Davis and Johnsrude, 2007; Huyck and Johnsrude, 2012; Cooke et al., 2022) - have shown similar results to the SWS studies described above, including a response similar to the PAN reported here when comparing responses to noise-vocoded speech after versus before training (Sohoglu et al., 2012; Sohoglu and Davis, 2016; Karunathilake et al., 2023). However, none of these studies used within-subject control stimuli [although (Karunathilake et al., 2023) used a control group similar to (Cheng et al., 2021) to rule out order effects], and all are potentially confounded by the task-relevance. Although similarly confounded, neuroimaging studies have consistently observed differential levels of activity in (particularly left) AC and adjacent regions for noise-vocoded speech when heard as speech (Scott et al., 2000, 2006; Davis and Johnsrude, 2003; Narain et al., 2003; Obleser and Kotz, 2010; Wild et al., 2012; Erb et al., 2013; Murai and Riquimaroux, 2021).

### Task correlates

By comparing neural responses between phases 2 and 3, we identified correlates of task-relevance in the absence of gross changes in perception (Fig. 4). Such task-related activity was observed as (i) a P3b (Polich, 2007; Halgren, 2008) from ∼400 – 800 ms, (ii) a late anterocentral negativity from ∼400 – 800 ms and (iii) longer and more generalizable decoding in phase 3 (SWS task-relevant) versus phase 2 (SWS task-irrelevant) (Fig. 5). However, we did not find significant ERP clusters for the interaction between phase (phase 3 versus 2) and stimulation type. This suggests that while the task introduced confounds (e.g. target detection, word identification, working memory, decision making), they did not strongly depend on perception of SWS as speech. This could reflect the fact that SWS and frequency-flipped SWS sound perceptually similar (especially compared to pure tones) and therefore were both treated as potentially task-relevant in phase 3. Another control for examining task-relevance that could mitigate effects of word identification (Koivisto et al., 2017) could be to use SWS tokens that are perceived as speech (as ours were) but that are not task-relevant, where we would expect the size of the P3b to co-vary with similarity of the foil stimuli to the targets (Pitts et al., 2014). In any case, the absence of the P3b in the task-irrelevant contrast (phase 2 versus 1) and its presence in the task-relevant contrast (phase 3 versus 2) is consistent with recent work in both visual and auditory perceptual awareness (Pitts et al., 2012; Cohen et al., 2020; Sergent et al., 2021).

## Conclusion

Using a three-phase paradigm in which we independently manipulated perception and task-relevance, we identified a novel neural difference linked with perceptual awareness of SWS words, which we interpreted as a speech-specific version of a perceptual awareness negativity (Dembski et al., 2021). Future work involving the paradigm developed here along with other methods may help reveal the precise neural mechanisms underlying conscious speech perception and help address diverging predictions from neuroscientific theories of consciousness (Seth and Bayne, 2022).

